# UDSMProt: Universal Deep Sequence Models for Protein Classification

**DOI:** 10.1101/704874

**Authors:** Nils Strodthoff, Patrick Wagner, Markus Wenzel, Wojciech Samek

## Abstract

**Motivation:** Inferring the properties of a protein from its amino acid sequence is one of the key problems in bioinformatics. Most state-of-the-art approaches for protein classification tasks are tailored to single classification tasks and rely on handcrafted features such as position-specific-scoring matrices from expensive database searches. We argue that this level of performance can be reached or even be surpassed by learning a task-agnostic representation once, using self-supervised language modeling, and transferring it to specific tasks by a simple finetuning step.

**Results:** We put forward a universal deep sequence model that is pretrained on unlabeled protein sequences from Swiss-Prot and finetuned on protein classification tasks. We apply it to three prototypical tasks, namely enzyme class prediction, gene ontology prediction and remote homology and fold detection. The proposed method performs on par with state-of-the-art algorithms that were tailored to these specific tasks or, for two out of three tasks, even outperforms them. These results stress the possibility of inferring protein properties from the sequence alone and, on more general grounds, the prospects of modern natural language processing methods in omics.

**Availability:** Source code is available under https://github.com/nstrodt/UDSMProt.

## 1 Introduction

Inferring protein properties from the underlying sequence of amino acids (primary structure) is a long-standing theme in bioinformatics and is of particular importance in the light of advances in sequencing technology and the vast number of proteins with mostly unknown properties. A rough estimate for this number is given by the size of the sparsely annotated TrEMBL dataset (158M) and should be set into perspective by comparison to the size of well-curated Swiss-Prot [The UniProt Consortium, 2018] dataset (560K) with much more complete annotation of protein properties.

There is a large body of literature on methods to infer protein properties, most of which make use of additional handcrafted features in addition to the primary sequence alone [Shen and Chou, 2007, Håndstad et al., 2007, Gong et al., 2016, Cozzetto et al., 2016, Li et al., 2017, 2018, Dalkiran et al., 2018]. These include experimentally determined functional annotations (such as Pfam [El-Gebali et al., 2018]) as well as features incorporating information from homologous (evolutionary related) proteins that are typically inferred from well-motivated but still heuristic methods such as the basic local alignment search tool (BLAST) [Madden, 2013], that searches a database for proteins that are homologous to a given query protein, via multiple sequence alignment. Handcrafted features that are based on experimental results rely on a preferably complete functional annotation and are therefore likely to fail to generalize for incompletely annotated proteins[Price et al., 2018]. Handcrafted features that are derived from multiple sequence alignments using alignment algorithms typically scale at least linearly with query and database size. This time complexity is not able to keep up with the present size and the exponential growth rates of present protein databases.

These bottlenecks urge for the development of methods that allow to directly predict protein properties from the sequence of amino acids alone, which is therefore a topic on the agenda of many research institutions [Bileschi et al., 2019, Rives et al., 2019, Rao et al., 2019]. Methods from deep learning, and self-supervised algorithms from natural language processing (NLP) in particular, are promising approaches in this direction.

Also the Machine Learning community recently gained interest in protein classification as possible application area for deep learning methods (see e.g. AlQuraishi [2019], Bileschi et al. [2019], Rives et al. [2019], zu Belzen et al. [2019], Rao et al. [2019]). In NLP, self-supervised approaches have shown tremendous prospects across a wide variety of tasks [Peters et al., 2018, Howard and Ruder, 2018, Radford et al., 2018, Devlin et al., 2018, Radford et al., 2019, Song et al., 2019, Yang et al., 2019, Liu et al., 2019], which rely on leveraging implicit knowledge from large unlabeled corpora by pretraining using language modeling or related unmasking tasks. This approach goes significantly beyond the use of pretrained word embeddings, where only the embedding layer is pretrained whereas the rest of the model is initialized randomly.

Protein classification tasks represent an tempting application domain for such techniques exploiting the analogy of amino acids as words and proteins and their domains as text paragraphs composed of sentences. In this setting, global protein classification tasks, such as enzyme class prediction, are analogous to text classification tasks (e.g. sentiment analysis). Protein annotation tasks, such as secondary structure or phosphorylation site prediction, map to text annotation tasks, such as part-of-speech tagging or named entity recognition. While this general analogy has been recognized and exploited already early on [Asgari and Mofrad, 2015], self-supervised pretraining is a rather new technique in this field [Rives et al., 2019, Rao et al., 2019]. Existing literature approaches in this direction [Rives et al., 2019, Rao et al., 2019] show significant improvements of models that were pretrained using self-supervision compared to their counterparts trained from scratch on a variety of tasks and demonstrate that models leverage biologically sensible information from pretraining. However, none of them explicitly demonstrated for these problems that pretraining can bridge the gap to state-of-the-art approaches that mostly rely on handcrafted features such as position-specific-scoring matrices (PSSM) derived via BLAST.

Our main contributions in this paper are the following: (1) We put forward a universal deep protein classification model for protein classification (UDSMProt) that is pretrained on Swiss-Prot and finetuned on specific classification tasks without any further task-specific modifications. (2) We demonstrate for three classification tasks that this model is able to reach or even surpass the performance level of state-of-the-art algorithms many of which make use of PSSM features and hence, to the best of our knowledge for the first time, the feasibility of inferring protein properties from the sequence alone. (3) We demonstrate the particular effectiveness of our approach for small datasets.

## 2 Algorithms and Training Procedures

### 2.1 UDSMProt: Universal Deep Sequence Models for Protein Classification

The idea of *UDSMProt* is to apply self-supervised pretraining to a state-of-the-art recurrent neural network architecture using a language modeling task. In this way the model learns implicit representations from unlabeled data that can be leveraged for downstream classification task. We aim to address a range of different classification problems within a single architecture that is universal in the sense that only the dimensionality of the output layer has to be adapted to the specific task, which facilitates the adaption of the approach to classification tasks beyond the three exemplary tasks considered in this work. For finetuning on the downstream classification tasks, all embedding weights and LSTM weights are initialized using the same set of weights obtained from language model pretraining. As we will demonstrate, this is a particularly powerful choice for small datasets.

Our proposed method relies on an *AWD-LSTM* language model [Merity et al., 2017], which is, at its heart, a 3-layer LSTM regularized by different kinds of dropouts (embedding dropout, input dropout, weight dropout, hidden state dropout, output layer dropout). During language model training, the output layer is tied to the weights of the embedding layer. Specific model parameters are listed in Table 6. The training procedure for transfer learning is largely inspired by *ULMFit* [Howard and Ruder, 2018] and proceeds as follows: In a first step, we train a language model on the Swiss-Prot database. In a second step, the language model’s output layer is replaced by a concat-pooling-layer [Howard and Ruder, 2018] and two fully connected layers, see Figure 1 for a schematic illustration. When finetuning the classifier, we gradually unfreeze layer group by layer group (four in total) for optimization, where we use discriminative learning rates i.e. we reduce the learning rate by a factor of 2 compared to the previous layer group [Howard and Ruder, 2018]. A single model is by construction only able to capture the context in a unidirectional manner, i.e. processing the input in the forward or backward direction. As simplest approach to incorporate bidirectional context into the final prediction, we train separate forward and backward models both for language models as well as for the finetuned classifiers. Finally, an ensemble model is obtained by averaging the output probabilities of both classifiers. We use a 1-cycle learning rate schedule [Smith, 2018] during training for 30 epochs during the final finetuning step. Any kind of hyperparameter optimization was performed based on the model performance on a separate validation set, while we report performance on a separate test set. Our way of addressing the specific challenges of the remote homology datasets are described in Section 3.4. In all cases, we use binary/categorical crossentropy as loss function and the AdamW optimizer [Loshchilov and Hutter, 2019].

**Figure 1:**
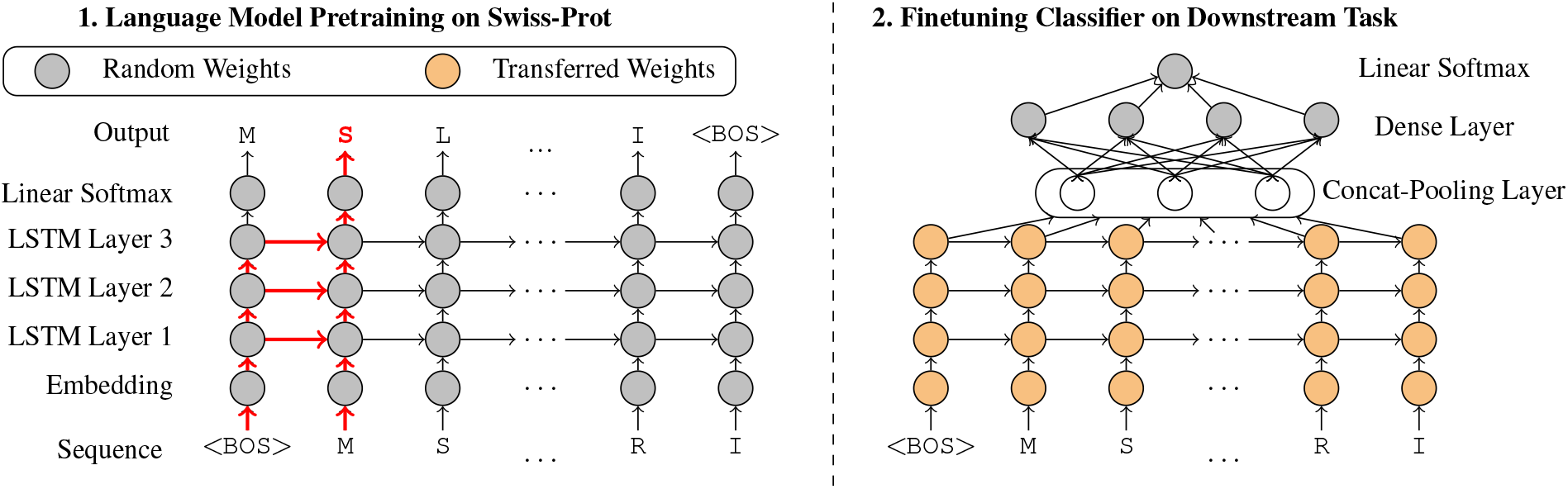
Schematic illustration of the training procedure, here for the amino acid sequence `<BOS>MSLR…RI`. The `<BOS>`-token marks the beginning of the sequence. The red arrows show the context for forward LM for predicting next character (S) given sequence `<BOS>M` of length 2. For finetuning on the downstream classification tasks, all embeddings weights and LSTM weights are initialized using the same set of weights obtained from language model pretraining. This has to be contrasted with the use of pretrained embeddings, where just the embedding weights are initialized in a structured way before the downstream finetuning step.

Note that a potential intermediate step where one finetunes the generic language model on the corpus underlying the classification step, as proposed by Howard and Ruder [2018], did only show an improvement in terms of language model quality but did not result in an improved downstream classification performance. This step was therefore omitted for the results presented below.

### 2.2 Baseline Model

In our experiments below, we mostly compare directly to reported results from approaches in the literature on predefined datasets. However, this does not allow for in-depth comparisons that modify for example details of the training procedure. To still allow to relate the results of the proposed method to state-of-the-art performance, we use a baseline model that reaches state-of-the-art performance on literature benchmarks and that can henceforth be used as proxy for models considered in the literature.

The performance of literature approaches on many protein classification tasks has been driven to a large extend by the inclusion of different kinds of handcrafted features rather than sophisticated model architectures or training procedures. The most beneficial input features throughout a variety of different classification tasks are obviously the position specific scoring matrices (PSSM) based on a multiple sequence alignment computed via position specific iterative BLAST (PSI-BLAST) [Madden, 2013]. PSI-BLAST is used to compare query sequences with a given database of already existing sequences, where the result is a list of local alignments solved with heuristics instead of using more time-consuming optimal local alignments with the Smith-Waterman-algorithm. PSI-BLAST is then used to find more distant relatives of a query protein, where a list of closely related proteins is created to get an initial general profile sequence. This profile sequence is used as a new query for the next iteration where a larger list of proteins is found for which again a profile sequence is computed. This process is repeated to a desired number of iterations. In our experiments we used the same parameters as reported in the literature [Li et al., 2018, Dalkiran et al., 2018, Shen and Chou, 2007], namely three iterations with `e_value` = 0.001, where `e_value` relates to the threshold for which an alignment is considered as significant. While the raw sequences from Swiss-Prot contained 26 unique amino acids (20 standard and 6 non-standard amino acids), PSSM features are computed only for the 20 standard amino acids. The raw sequence of length *L* was then one-hot encoded into an *L* × 26 matrix which is concatenated with the *L* × 20 PSSM feature matrix yielding an *L* × 46 input matrix overall. To make use of the full parallelization capabilities while retaining most information at the same time, we padded the sequences to a maximum length of 1024 residues.

For all following experiments we used a Convolutional Neural Network (CNN) consisting of seven layers, where each convolutional layer was followed by rectified linear unit (ReLU) and max pooling by a factor of 2. The number of filters across layers are: 1024, 512, 512, 512, 256, 256, 256 (with valid padding mode) each with filter size of 3, after the last layer the dimensionality was 6 × 256. The convolutional stage was followed by flatten/vectorization and three dense layers (512, 256, 128) each followed by dropout (with 25% dropout rate) and finally a softmax layer with six nodes, one for each class. For all models we minimized either binary or categorical crossentropy (for level 0 and level 1 respectively) with AdaMax, which is a variant of adaptive moment estimation (Adam) based on the infinity norm [Kingma and Ba, 2015]. The hyperparameters follow those provided in the paper (learning rate 0.002, exponential decay rates *β*_1_ = 0.9, *β*_2_ = 0.999).

## 3 Results and Discussion

The results are organized as follows: We start by discussing language modeling as baseline task in Section 3.1, before demonstrating the capabilities of *UDSMProt* on three prototypical protein classification tasks, namely enzyme class prediction in section 3.2, gene ontology prediction in section 3.3 and remote homology detection in section 3.4. While for the latter two we compare to literature in a straightforward manner, for enzyme class prediction we provide a more extensive evaluation highlighting several important aspects.

### 3.1 Language Modeling

The language modeling task involves predicting the next token for a given sequences of tokens and is one of the key NLP tasks for demonstrating the general understanding of a language. In this particular case it evaluates the implicit knowledge about the structure of proteins that has been captured by the model, which can potentially be leveraged for downstream classification tasks.

Our language model operates on protein sequence data tokenized on the level of amino acids (i.e. character-based, see fig. 1). Interestingly, the language model performance depends strongly on the way the similarity threshold is incorporated in the train-test split procedure. For this reason, we split the data into train, validation and test set with ratios 90%:5%:5%, where we compare two methods: (1) randomly, i.e. without taking sequence similarity into account and (2) based on UniRef50 cluster assignments. While the first model reaches a perplexity of 6.88 and a corresponding prediction accuracy of 0.409, the second model only reaches a perplexity of 11.75 with 0.244 accuracy. However, these differences in language model performance do not lead to measurable differences in the downstream performance, see Appendix 3 for a detailed discussion. In conclusion this experiment demonstrates a rather profiencient knowledge of the respective language model of the underlying construction principles for proteins.

### 3.2 Enzyme Class Prediction

We start our analysis on downstream classification tasks with EC classification for the reason that it is a conceptually simple task for which a large number of annotated examples is available. The experiments in this section are organized as follows: Section 3.2.2 presents an in-depth analysis of the effects of sequence similarity and redundant sequences for the proposed *UDSMProt* in comparison to the baseline model operating on PSSM features. Section 3.2.3 compares the performance of both models to results reported in the literature and establishes the baseline model as a proxy for state-of-the-art approaches from the literature. Finally, in Section 3.2.4 we investigate for both models the dependence on the size of the training dataset by artificially reducing its size. This allows to illustrate the particular advantages of the proposed *UDSMProt* in the regime of small datasets.

#### 3.2.1 Task and Datasets

Enzyme prediction is a functional prediction task targeted to predict the Enzyme Commission number. The enzyme commission number is a hierarchical numerical classification scheme for enzymes based on the chemical reactions they catalyze. In particular, we consider EC prediction for level 0, i.e. predicting enzyme vs. non-enzyme, and level 1, i.e. predicting one of the six main enzyme classes. A powerful EC classification algorithm of the pre-deep-learning-era was provided by *EzyPred* [Shen and Chou, 2007], which owed its success to the design of a hierarchical approach and to appropriate input features which are a combination of the functional (BLAST against a PFAM database) and evolutionary information (PSI-BLAST [Madden, 2013] against the Swiss-Prot database). For hierarchical classification (level 0 to level 2), a simple k-nearest-neighbor classifier (KNN) was trained in order to achieve convincing results. *EzyPred* was superseded by *DEEPre* [Li et al., 2018] where deep learning was applied to raw sequence and homology data as input. Instead of training simple classifiers on highly engineered features, they trained feature representation and classification in an end-to-end fashion with a hybrid CNN-LSTM-approach. Recently, *ECPred* [Dalkiran et al., 2018] also showed competitive results by building an ensemble of well-performing classifiers (Subsequence Profile Map with PSSM [Sarac et al., 2008], BLAST-kNN [Madden, 2013] and Pepstats-SVM using peptides statistics [Rice et al., 2000]). Nevertheless, drawbacks as described in Section 1 remain, i.e. requiring functional annotations of homologous proteins, which is not guaranteed for evolutionary distant or insufficient annotated proteins.

In addition to the existing DEEPre (similarity threshold 40%) and ECPred (similarity threshold 50%) datasets [Li et al., 2018, Dalkiran et al., 2018] that provide only representative sequences, we also work with two custom datasets EC40 and EC50 (similarity threshold 40% and 50%) by combining best practices for the dataset construction, see Appendix 1 for a detailed description.

#### 3.2.2 Effect of Similarity Threshold and Redundant Sequences

In order to investigate the benefits of the proposed approach in comparison to algorithms relying on alignment features, we based our initial analysis on the custom EC40 and EC50 datasets, which are constructed analog to datasets in the literature. This approach represents a very controlled experimental setup, where one can investigate the effect of the chosen similarity threshold, the impact of redundant sequences during training and potential sources of data leakage during pretraining in a reliable way.

We base our detailed analysis of the proposed method *UDSMProt* compared to a baseline algorithm operating on PSSM features on EC prediction tasks at level 0 (enzyme vs. non-enzyme) and level 1 (main enzyme class). In particular, we aim to investigate the impact of non-redundant sequences when training the baseline classifier and the impact of different similarity thresholds. It is a well-known effect that the difficulty of the classification problem scales inversely with the similarity threshold, as a higher similarity threshold leads to sequences in the test set that are potentially more similar to those seen during training. In the extreme case of a random split, i.e. by disregarding cluster information, the test set performance merely reflects the algorithm’s capability to approximate the training set rather than the generalization performance when applied to unseen data. The failure to correctly incorporate the similarity threshold is one of the major pitfalls for newcomers in the field. Here, we perform level 0 and level 1 prediction on two different datasets, namely EC40 (40%) and EC50 (50% similarity cutoff). Both datasets only differ in the similarity thresholds and the version of the underlying Swiss-Prot databases.

If not noted otherwise, CNN models are trained on representative sequences as this considerably reduces the computational burden for determining PSSM features and is in line with the literature, see e.g. [Li et al., 2018, Dalkiran et al., 2018], whereas *UDSMProt* is conventionally trained using the full training set including redundant sequences, whereas the corresponding test and validation sets always contain only non-redundant sequences. For the EC50 dataset non-redundant sequences enlarge the size of the training set from 45k to 114k and from 86k to 170k sequences for level 1 and level 0 respectively. For EC40 the size is enlarged from 20k to 100k and from 46k to 150k for level 1 and level 0 respectively.

In Table 1, we compare the two classification algorithms *UDSMProt* and the baseline CNN that were introduced in Section 2 in terms of classification accuracy, which is the default metric considered in the literature for this task. There is a noticeable gap in performance across all experiments between CNN(seq; non-red.) and CNN(seq+PSSM; non-red.) which is a strong indication for the power of PSSM features. This gap can be reduced by the use of redundant sequences from training clusters (CNN(seq)) but still remains sizable. Most importantly, however, the gap can be closed by the use of language model pretraining. Disregarding the case of the EC40 dataset at level 0, the best-performing *UDSMProt* outperforms the baseline algorithms that make use of PSSM features. Combining information from both forward and backward context consistently improves over models with unidirectional context. As another observation, pretraining leads to a consistent advantage compared to models trained from scratch that cannot be compensated by increasing the number of training epochs for the models trained from scratch.

**Table 1.** EC classification accuracy on the custom EC40 and EC50 datasets. Here and throughout the paper we use the following abbreviations: fwd/bwd (training in forward/backward direction), seq (raw sequence as input), non-red. (training on non-redundant sequences i.e. representatives only), pretr. (using language model pretraining), leak. (leakage;PSSM features computed on the full dataset). The best performing classifiers are marked in bold face.

|  |  | EC40 |  | EC50 |  |
| --- | --- | --- | --- | --- | --- |
| Method |  | Level 0 | Level 1 | Level 0 | Level 1 |
| Baseline<sup>†</sup> | seq;non-red. | 0.83 | 0.38 | 0.88 | 0.71 |
|  | seq | 0.84 | 0.61 | 0.92 | 0.80 |
|  | seq+PSSM;non-red.;clean | 0.91 | 0.84 | 0.95 | 0.94 |
|  | seq+PSSM;non-red.;leak. | **0.92** | 0.85 | 0.95 | 0.95 |
| UDSMP<sub>Prot</sub><sup>†</sup> | fwd;pretr.; non-red. | 0.82 | 0.79 | 0.93 | 0.94 |
|  | fwd;from scratch | 0.87 | 0.79 | 0.94 | 0.94 |
|  | fwd;pretr. | 0.89 | 0.84 | 0.95 | 0.96 |
|  | bwd;pretr. | 0.90 | 0.85 | 0.95 | 0.96 |
|  | fwd+bwd;pretr. | 0.91 | **0.87** | **0.96** | **0.97** |

Finally, Table 1 illustrates that the *UDSMProt* classification models benefit from redundant training sequences for the downstream classification task, where the benefit is greater as the similarity threshold decreases. Comparing corresponding results from different similarity thresholds, i.e. results from EC40 to those from EC50, reveals the expected pattern, in the sense that lowering the similarity threshold complicates the classification task as test sequences show smaller overlap with sequences from the training set.

Pretraining or precomputing features such as PSSMs on the full dataset disregarding the train-test splits for the downstream classification tasks is a possible source of data leakage that can lead to a systematic over-estimation of the model’s generalization performance by implicitly leveraging information about the test set during the training phase. Here, we investigate the issue for the case of PSSM features as this issue was, to the best of our knowledge, not addressed in the literature so far. To this end, we compute two sets of PSSM features, one set computed based on the whole Swiss-Prot database (corresponding classification model: CNN(seq+PSSM; non-red.; leakage)) and a separate set based only on cluster members from the training data (corresponding classification model: CNN(seq+PSSM; non-red.; clean)). It turns out that the model with PSSM features computed on a consistent train-test-split always performs slightly worse than its counterpart that relies on PSSM features computed on the whole dataset. However, from a practical perspective, the effect of test data leakage remains small. In Appendix 5, we provide a more extensive evaluation of the effect by varying the size of the training database that is used for calculating PSSM features.

To reiterate the main findings of the experiments carried out in this section, the most crucial observation is that language model pretraining is capable of closing the gap in performance between models operating on PSSM features compared to models operating on the sequences alone. The second main observation is that redundant sequences rather than cluster representatives only have a positive impact on the downstream classification training. The most obvious explanations for this observation are inhomogeneous clusters that contain samples with different labels that carry more finegrained information than a single label per cluster representative. Finally, the effect of data leakage from computing PSSM features on the whole dataset disregarding train-test splits turned out to be small, but should be kept in mind for future investigations.

#### 3.2.3 Comparison to Literature Benchmarks

In order to relate our proposed approach to state-of-the-art methods in literature, we conducted an experiment on two datasets provided by ECPred[Dalkiran et al., 2018] and *DEEPre[Li et al., 2018]*. One of the purposes of this analysis is to justify our choice of the CNN baseline algorithm by demonstrating that it performs on par with state-of-the-art algorithms that do not make use of additional side-information e.g. in the form of PFAM features. When comparing to literature results on the *DEEPre* dataset, we exclude models relying on Pfam features from our comparison. Leaving aside the very unfavorable scaling with the dataset size [Bileschi et al., 2019] and possible issues with data leakage due to features computed on the full dataset, methods relying on these features will fail when applied to proteins without functional annotations, see also the discussion in [Dalkiran et al., 2018]. In fact, a recent study estimated that at least one-third of microbial proteins can not be annotated through alignments on given sequences [Price et al., 2018]. Most notably, this excludes the most elaborate *DEEPre* [Li et al., 2018] model (with 0.96 level 0 and 0.95 level 1 accuracy on the *DEEPre* dataset) and *EzyPred* [Shen and Chou, 2007] (with 0.91 level 0 and 0.90 level 1 accuracy) from the comparison.

Table 2 shows the results of this experiment, see Appendix 4 for details on the evaluation procedure. Note, that a convolutional model (as our baseline) seemed sufficient when compared to the hybrid model of *DEEPre* (using convolutional layers followed by a recurrent layer (LSTM)) as can been seen in Table 2 where our baseline even surpassed the reported performances (91% vs. 88% for level 0 and 84% vs. 82% for level 1). Also for testing on *ECPred* our baseline approach yielded competitive results indicating a well-chosen baseline model. These results justify a posteriori our design choices for the CNN baseline model.

**Table 2.**
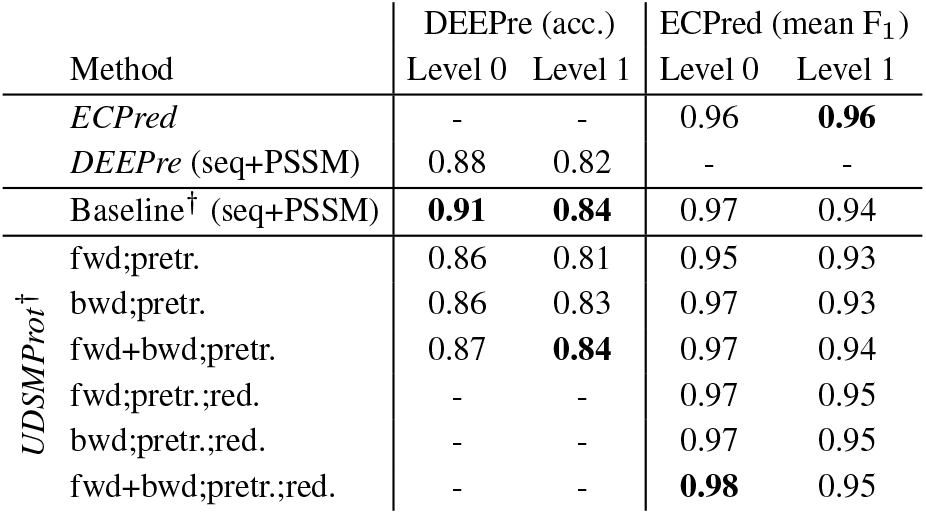
EC classification accuracy on the published DEEPre and ECPred datasets compared to literature results from DEEPre [Li et al., 2018], and ECPred [Dalkiran et al., 2018] disregarding models relying on Pfam features. Results on the DEEPre dataset were evaluated using 5-fold crossvalidation. Results established in this work are marked by ^†^.

Turning to the performance of the proposed *UDSMProt*, we find a solid prediction performance reaching state-of-the-art performance reported in the literature for algorithms operating on PSSM features. Considering the results of the previous section, the results on the *DEEPre* dataset represent only the lower bound for the achievable performance as it profits considerably from redundant training sequences, which could, however, not be reconstructed from the given representatives without the underlying cluster assignments. Considering the sizable performance gaps between training on redundant and non-redundant datasets in Table 1, it is even more remarkable that *UDSMProt* already reaches state-of-the-art performance when trained on non-redundant sequences. For *ECPred* we report both the performance for training on the original training set as well as the performance on a redundant training set comprising all corresponding Uniref50 cluster members as shown in the three bottom lines in Table 2. In terms of level 0 performance the proposed approach outperforms *ECPred* and it shows competitive performance at level 1.

To summarize, our baseline model reaches state-of-the-art performance compared to literature approaches disregarding those that incorporate features from functional annotations (such as PFAM) and can therefore be used as proxy for state-of-the-art algorithms in the following investigations. This finding enhances a posteriori also the significance of the results established for the EC40 and EC50 datasets in Section 3.2.2. The proposed *UDSMProt*-model is very competitive on both literature datasets.

#### 3.2.4 Impact of Dataset Size

In this section, we aim to demonstrate the particular advantages of the proposed *UDSMProt*-approach in the regime of small dataset sizes. To investigate this effect in a clean experimental setup, we conducted an experiment with consecutively decreasing training set sizes, while keeping test and validation sets fixed. The hyperparameters were kept fixed to those of the run with full training data.

For this experiment, we used the EC50 dataset as described in the appendix with numbers per class as shown in Table 5 and trained a level 1 classifier for each split. We compared our proposed approach (AWD-LSTM with pretraining and from scratch) with baseline models (CNN with PSSM features and CNN on redundant sequences only) for seven different training set sizes measured in terms of clusters compared to the number of clusters in the original training set.

The results from Figure 2 show an interesting pattern: The bidirectional *UDSMProt* model always outperforms the CNN baseline model and most interestingly the gap between the two models increases for small dataset sizes, which suggests the representations learned during language model finetuning represent a more effective baseline for finetuning than using PSSMs as fixed input features. As a second observation, also the gap to the models trained from scratch widens. Reducing the number of training clusters by 50% only leads to a decrease in model performance by 3%, whereas the performance of the model trained from scratch drops by 8%.

**Figure 2:**
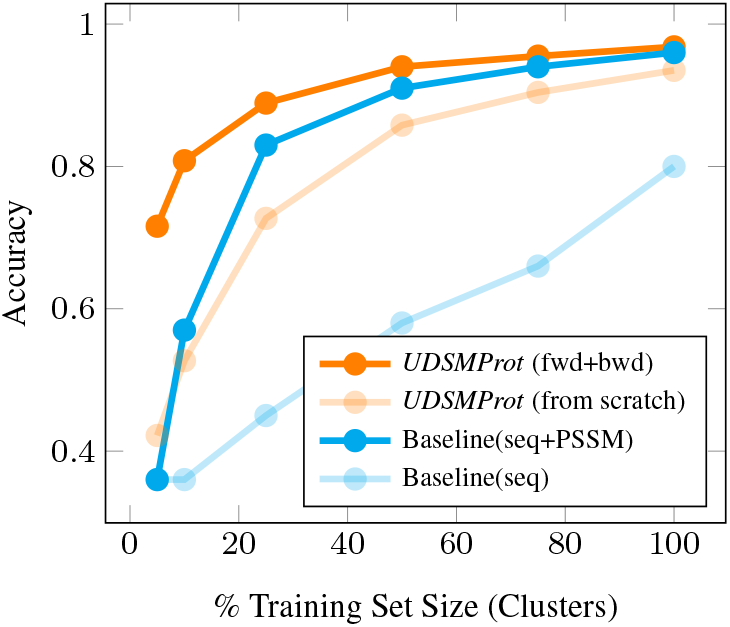
Dependence of the EC classification accuracy (level 1; EC50 dataset) on the size of the training dataset. *UDSMProt* performs better than the baseline model also in the regime of small datasets that is particularly important for practical applications.

To summarize, both observations from above represent strong arguments for applications of *UDSMProt* in particular to small datasets. Our results suggest to make language model pretraining a standard procedure in these cases.

### 3.3 Gene Ontology Prediction

To stress the universality of the proposed approach, we also present results for gene ontology prediction, which is a functional multi-label classification task.

#### 3.3.1 Task and Dataset

A more general although closely related problem to enzyme prediction is gene ontology prediction. Gene ontology is an international bioinformatics initiative to unify a part of the vocabulary for representation of proteins attributes. It covers three domains, namely cellular components, molecular functions and biological processes. The nomenclature is organized into hierarchies ranging from coarse to fine-grained attributes. Here, we focus on the domain of molecular function prediction. Similar to enzyme class prediction, the first proposed approaches in this field relied on handcrafted features like functionally discriminating residues (FDR) with PSSM [Gong et al., 2016] and classification models consisting of an array of Support Vector Machines [Cozzetto et al., 2016]. Recently, deep learning approaches have raised the bar by using convolutional neural networks [Kulmanov et al., 2017] and residual neural networks [zu Belzen et al., 2019].

For convienient comparability with literature results, we focus on protein annotations belonging to the molecular function (MF) and completely disregard the other two main categories, cellular component and biological process. This is also the considered case of *DeeProtein* [zu Belzen et al., 2019], with data from the CAFA3 challenge [Radivojac et al., 2013], which is a major attempt to establish a suitable evaluation framework for protein function prediction. We use the scripts provided by zu Belzen et al. [2019] to construct appropriate training and test sets. Specifically, we train on their Swiss-Prot training set, that was designed to have sequence similarities below 50% compared to the CAFA3 test set, resulting in a training set of 135468 unique sequences. In order to be able to compare directly to literature results [Kulmanov et al., 2017, zu Belzen et al., 2019], we compare performance on a CAFA3 benchmark set comprising 575 sequences for a selection of 539 molecular function GO-terms that were predicted by *DeepGO*.

#### 3.3.2 Experimental Setup and Results

We used the implementation provided by zu Belzen et al. [2019] to construct train and test datasets. We grouped duplicate sequences across multiple species with slightly different sets of annotations by building a grand union on those in order to ensure that every possible annotations is considered. Also the evaluation of the final scores from the raw model predictions was carried out using the provided evaluation scripts [Kulmanov et al., 2017, zu Belzen et al., 2019]. The reported metrics are based on those reported in the CAFA3 challenge, namely a protein-centric maximum F-measure F_max_ along with a corresponding average recall and a term-centric AUC [Clark and Radivojac, 2013, Kulmanov et al., 2017].

The results in Table 3 show that *UDSMProt* model outperforms the existing approaches in terms of F_max_-score and recall by a large margin and perform on par in terms of AUC. As for EC prediction, the best-performing *UDSMProt*-model is the forward-backward-ensemble model that exploits bidirectional context. The result presented in this section substantiate our claims regarding the universality of transferring implicit knowledge to task-specific requirements.

**Table 3.** Gene ontology (molecular function) prediction performance evaluated on the DeepGO [Kulmanov et al., 2017] benchmark set in comparison to literature results from DeeProtein [zu Belzen et al., 2019], DeepGO [Kulmanov et al., 2017], FFPred3 [Cozzetto et al., 2016], and GoFDR [Gong et al., 2016].

| Method | AUC | F<sub>max</sub> | Recall |
| --- | --- | --- | --- |
| *DeeProtein* (Swiss-prot) | 0.89 | 0.51 | 0.47 |
| *DeeProtein* (CAFA3) | 0.88 | 0.50 | 0.42 |
| *DeepGO* | **0.90** | 0.48 | 0.40 |
| *FFPred3* | 0.86 | 0.38 | 0.40 |
| *GoFDR* | 0.84 | 0.52 | 0.36 |
| *UDSMProt*<sup>†</sup> fwd;from scratch | 0.88 | 0.57 | 0.50 |
| fwd;pretr. | 0.89 | 0.59 | 0.53 |
| bwd;pretr. | 0.88 | 0.59 | 0.54 |
| fwd+bwd;pretr. | 0.89 | **0.60** | **0.54** |

### 3.4 Remote Homology and Fold Detection

As third demonstration of the universality of our approach, we consider remote homology detection tasks. The corresponding datasets consist of a few hundred training examples and are thus situated clearly in the small dataset regime investigated in Section 3.2. This substantiate the claims made in Section 3.2 in a real-world setting.

#### 3.4.1 Task and Datasets

Remote homology detection is one of the key problems in computational biology and refers to the classification of proteins into structural and functional classes, which is considered to be a key step for further functional and structural classification tasks. Here, we consider remote homology detection in terms of the SCOP database [Murzin et al., 1995], where all proteins are organized in four levels: class, fold, superfamily and family. Proteins in the same superfamily are homologous and proteins in the same superfamily but in different families are considered to be remotely homologous. Remote homology detection has a rich history and the interested reader is referred to a review article on this topic by Chen et al. [2016]. We will compare to *ProDec-BLSTM* [Li et al., 2017] with a bidirectional recurrent neural network operating on PSSM input features building on earlier work [Hochreiter et al., 2007]. A classical baseline method is provided by *GPkernel* [Håndstad et al., 2007], who apply kernel-methods to sequence motifs.

For remote homology detection, we make use of the SCOP 1.67 dataset as prepared by Hochreiter et al. [2007], which has become a standard benchmark dataset in the field. Here, the problem is framed as a binary classification problem where one has to decide if a given protein is contained in the same superfamily or fold as a given reference protein. The superfamily/fold benchmark is composed of 102/85 separate datasets and we report the mean performance of all models across the whole set. The standard metrics considered in this context are AUC and AUC50, where the latter corresponds to the (normalized) partial area under the ROC curve integrated up to the first 50 false positives, which allows for a better characterization of the classifier in the domain of small false positive rates that is most relevant for practical applications than the overall discriminative power of the classifier as quantified by AUC.

#### 3.4.2 Experimental Setup and Results

The remote homology and fold detection tasks are challenging for two reasons: The datasets are rather small and the task comprises 102 or respectively 85 different datasets that would in principle require a separate set of hyperparameters. To keep the process as simple as possible, we decided to keep a global set of hyperparameters for all datasets of a given task. The procedure is as follows: As no validation is provided for the original datasets, we split the training data into a training and a validation set based on CD-HIT clusters (threshold 0.5). We optimize hyperparameters using the mean AUC for all datasets of a given task measured on the validation set. Most importantly, this involves fixing a (in this case constant) learning rate that is appropriate across all datasets. Using these hyperparameter settings, we perform model selection based on the validation set AUC, i.e. for each individual dataset, we select the model at the epoch with the highest validation AUC. We evaluate the test set AUC for these models and report the mean test set metrics.

The results of these experiments are shown in Table 4. Both for homology and fold detection according to most metrics, the *UDSMProt* model trained from scratch performs worse than the original LSTM model [Hochreiter et al., 2007]. This is most likely due to the fact that the *UDSMProt* model is considerably larger than the latter model and most datasets are fairly small with a few hundreds training examples per dataset. This deficiency is overcome with the use of language model pretraining, where both unidirectional models perform better than the LSTM baseline model. This observation is in line with the experiments in Section 3.2.4 that demonstrates the particular effectiveness of the proposed approach for small datasets. The best-performing model from the literature, *ProDec-BLSTM*, is a bidirectional LSTM operating on sequence as well as PSSM features. Interestingly, reaching its performance in terms of overall AUC required the inclusion of bidirectional context i.e. the forward-backward ensemble model. The proposed method also clearly outperforms classical methods such as *GPkernel* [Håndstad et al., 2007] both on the fold and the superfamily level. The excellent results on remote homology and fold detection support our claims on the universality of the approach as well as the particular advantages in the regime of small dataset sizes.

**Table 4.**
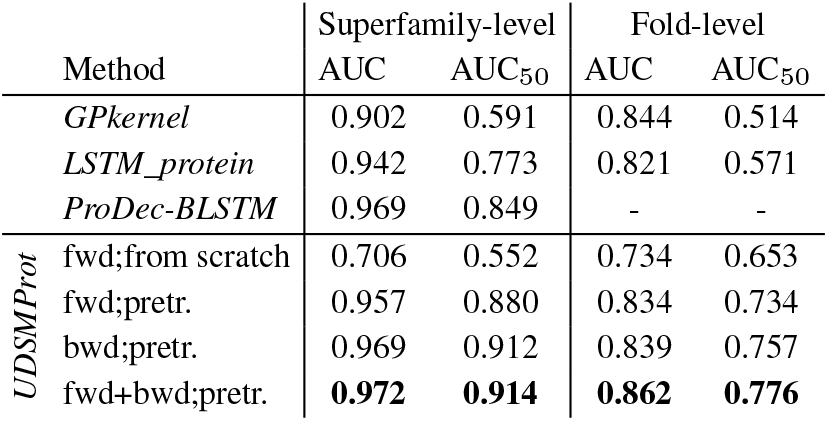
Remote homology and fold detection performance on the SCOP 1.67 benchmark dataset compared to literature results from GPkernel [Håndstad et al., 2007], LSTM_protein [Hochreiter et al., 2007], and ProDec-BLSTM [Li et al., 2017].

## 4 Summary and Outlook

In this work, we investigated the prospects of self-supervised pretraining for protein classification tasks leveraging the recent advances in NLP in this direction. Protein classification represents an ideal test bed for NLP methods. Most importantly, a single, universal model architecture with no task-specific modifications apart from a finetuning step that operates on the sequence of amino acids alone is able to reach or even exceed state-of-the-art performance on a number of protein classification tasks. This is achieved by powerful, implicitly learned representations from selfsupervised pretraining, whereas most state-of-the-art algorithms make use of PSSM features obtained from BLAST database searches that scale unfavorably with dataset size. In addition, the proposed method shows particular advantages for small datasets. Differently from typical NLP tasks, the dataset creation and the evaluation procedure has to be carried out with particular care, as small differences in the procedure can have large impact on the difficulty of the classification problem and hence on the comparability of different approaches. This applies in particular to a well-defined way of handling the similarity threshold, i.e. dealing with homologous sequences that differ only by a few amino acids when splitting into train and test sets. These factors urge for the creation of appropriate benchmark datasets that convert raw data from an exemplary subset of the many existing protein classification tasks into benchmark datasets in a transparent manner that allow for a rigorous testing of machine learning algorithms in this setting.

Given the insights gained from the three classification tasks, we can draw the following general conclusions for generic protein classification tasks:

1. Considering the fact that *UDSMProt* was able to reach or surpass state-of-the-art performance suggests that problem-specific architectures are less important than the training procedure, at least for models that are powerful enough. This allows to design task-independent, universal classification algorithms that can be applied without much manual intervention to unseen classification tasks.
2. Bidirectional context is important, which is reflected by the fact that in all cases forward-backward-ensemble reached the best performance and in most cases improved the performance of unidirectional models considerably. Ensembling forward and backward models is in fact the simplest – although at the same time a quite ineffective – way of capturing bidirectional context. From our perspective, this represents an opportunity for approaches such as BERT [Devlin et al., 2018, Liu et al., 2019] or XLNet [Yang et al., 2019], which are able to capture the bidirectional context directly. This might be particularly important for more complicated protein classification tasks that go beyond the prediction of a single global label such as sequence annotation tasks like secondary structure or phosphorylation site prediction.
3. Redundant sequences are a valuable source of information also for downstream classification tasks. This fact is in tension with the standard practice in bioinformatics, where in many cases only representatives without the corresponding cluster assignments are presented. To ensure comparability, benchmarking datasets should always include full information to reproduce the cluster assignments used during dataset creation, which would allow at least retrospectively to reconstruct the complete dataset from a given set of representatives.
4. Data leakage arising from inconsistent train-test splits between pretraining and classification e.g. by precomputing features (such as PSSM or Pfam features) on the full Swiss-Prot database without excluding downstream test clusters is a possible source of systematic overestimation of the model’s generalization performance. From our experiments on PSSM features in the context of EC prediction, its effect was found to be small in particular for large pretraining datasets such as Swiss-Prot, but it should be kept in mind for future investigations.

Leveraging large amount of unlabeled data in the form of large, in parts very well-curated protein databases by the use of modern NLP methods, represents a new paradigm in the domain of proteomics. It will be interesting to see how this process continues with the rapidly evolving algorithmic advances in the field of NLP, considering in particular approaches like BERT [Devlin et al., 2018, Liu et al., 2019], see also [Rives et al., 2019] for first applications in the domain of proteomics, and XLNet [Yang et al., 2019]. This is particularly relevant for more complicated protein classification tasks that go beyond the prediction of a single global label, such as sequence annotation tasks like secondary structure or phosphorylation site prediction, which even more require bidirectional contexts that cannot be captured by traditional language models.

## Acknowledgements

The authors thank J. Vielhaben for discussions and work on related topics. This work was supported by the Bundesministerium für Bildung und Forschung (BMBF) through the Berlin Big Data Center under Grant 01IS14013A and the Berlin Center for Machine Learning under Grant 01IS18037I. *UDSMProt* was implemented using Pytorch [Paszke et al., 2017] and fast.ai [Howard et al., 2018] and CNN baseline models were implemented in Keras [Chollet et al., 2015].

## 1 Custom Enzyme Class Datasets EC40 and EC50

As discussed in the main text, in addition to the original *DEEPre* and *ECPred* datasets, we recreated datasets for similarity threshold 40% and 50% by combining best practices from both [Li et al., 2018] and [Dalkiran et al., 2018]. The corresponding dataset with similarity threshold 40% relying on clusters from a prior CD-HIT [Fu et al., 2012] run is termed EC40. Compared to *DEEPre* we slightly adapted their dataset generation procedure by clustering Swiss-Prot all at once with CD-HIT, instead of clustering enzymes and non-enzymes separately, as this mimics the way UniRef50 clusters are constructed, albeit for similarity threshold 40% in this case. The dataset using a similarity threshold of 50% by using UniRef50-clusters is termed EC50. In both cases, we do not balance the number of enzymes and non-enzymes and restrict ourselves to proteins with a single EC-label.

Table 5 summarizes all datasets used for our experiments, namely the original data from [Li et al., 2018] and [Dalkiran et al., 2018] in the first two columns, and our replica in the last two columns.

**Table 5.** EC Prediction Datasets Overview

| EC class | DEEPre | ECPred | EC40 | EC50 |
| --- | --- | --- | --- | --- |
| Oxidoreductase | 3341 | 8242 | 3781 | 8244 |
| Transferase | 8517 | 20133 | 9137 | 20139 |
| Hydrolase | 5917 | 16018 | 7987 | 16020 |
| Lyase | 1532 | 3475 | 1401 | 3475 |
| Isomerase | 1193 | 2883 | 1160 | 2834 |
| Ligase | 1666 | 4429 | 2165 | 4429 |
| Enzymes | 22166 | 55180 | 25631 | 55141 |
| Non-Enzymes | 22142 | 25333 | 32387 | 49799 |
| Total | 44336 | 80513 | 58018 | 104940 |

The EC50 and EC40 EC classification datasets are constructed according to the following procedure:

1. Acquire Swiss-Prot (2016_08 for EC40 and 2017_03 for EC50).
2. Cluster Swiss-Prot with CD-HIT (40% similarity cutoff) for EC40 or acquire UniRef50 (2017_03) for EC50 and apply it to Swiss-Prot.
3. Remove non-enzymes which have not enough experimental evidence in order to avoid misleading false negatives (annotated evidence is greater or equal to 4).
4. Remove proteins which are annotated as fragments.
5. Remove enzymes with multiple enzymatic annotations in order to obtain a single-label classification problem.
6. Filter proteins by length to include only proteins with more than 50 and less than 5000 amino acids.
7. Split by clusters into 80% training and 10% validation and 10% test set.
8. For test and validation set, we select only representatives (from CD-HIT or UniRef50 clustering) or alternatively the first member of the cluster in the case where the representative was excluded by filtering rules. Optionally a similar reduction is applied to the training set to obtain a set of non-redundant sequences.
9. Use the predefined cluster assignments of the previous step and similarly distribute all remaining Swiss-Prot clusters onto training, validation and test dataset according to split ratios 90%, 5% and 5%, respectively, to obtain a clean train-test-split on the whole Swiss-Prot dataset consistent with chosen clusters for the downstream classification task. Again, we keep only representatives for validation and test set.

The last step is important for suitable database creation for PSI-BLAST for the experiments in Section 3.2.2 and Appendix 5. One thing can already be anticipated, as a result, for significantly more sequences from the test dataset PSI-BLAST returned empty queries resulting in less informative features for the test set.

## 2 AWD-LSTM Parameters

Table 6 lists the AWD-LSTM hyperparameters used for language modeling and classifier finetuning.

**Table 6.** AWD-LSTM Parameters

| Parameter | Value |
| --- | --- |
| Joint parameters |  |
| Number of hidden units | 1150 |
| Number of layers | 3 |
| Embedding dimension | 400 |
| Backpropagation through time (bptt) | 70 |
| Gradient clipping | 0.25 |
| Weight decay | 1e-7 |
| Language-model-specific parameters |  |
| Dropout ($p_o, p_h, p_i, p_e, p_w$) | $0.5 \cdot (0.25, 0.1, 0.2, 0.02, 0.15)$ |
| Classifier-specific parameters |  |
| Dropout ($p_o, p_h, p_i, p_e, p_w$) | $0.5 \cdot (0.4, 0.2, 0.6, 0.1, 0.5)$ |
| Max. length (explicit backprop.) | 1024 |
| Number of hidden units (head) | 50 |

## 3 Detailed Discussion of Language Model Performance

We present language model results using *AWD-LSTM* language models trained on Swiss-Prot using 90% / 5% / 5% splits based on clusters for training, validation and test set, respectively. We train using redundant sequences and evaluate on a reduced dataset with a single representative sequence for each cluster. As performance metrics, we report the perplexity (as natural exponential of the test loss) and the prediction accuracy. We tokenize on an amino acid level, the resulting vocabulary comprises the following additional tokens in addition to the 20+2 canonical amino acids: `X` (unknown), `B` (either `D` or `N`), `Z` (either `E` or `Q`), `<BOS>` (marks beginning of new protein). We stress that we do not specifically aim to optimize the language model performance, which could be easily improved using appropriate postprocessing techniques [Grave et al., 2016], as it only serves as pretraining objective in this context.

At this point a comment on the different training dataset sizes is appropriate. For random splits we disregard any cluster information and distribute samples according to the ratios 90% / 5% / 5%, which obviously results in the largest training dataset. In cases where cluster information is used the splits are performed by clusters, where we sort the clusters by the number of members and distribute them onto the three sets consecutively. Finally, we also report results for a dataset, where the train-test-split used for the language model respects those of a chosen downstream classification task, in this case level 1 EC prediction on the EC50 dataset described below. This construction avoids potential data leakage from using implicit knowledge about test and validation sets that is contained in the language model representations in the actual downstream classification task, see Section 3.2.2 for a detailed discussion.

The results on language modeling are compiled in Table 7. As expected, the best language model is reached on random dataset splits with perplexities of around 7. It is the coverage of the whole dataset in terms of included clusters that is most important for language model performance. This is apparent from the comparison of (1) the language model trained on a random split, where we artificially restricted the training set to 64% of the original training set compared to (2) the UniRef50 runs. In both cases, the size of the training set is comparable but the former (1) reaches a perplexity comparable to that of the model trained on the full dataset with random splits, whereas the latter (2) only reaches a perplexity of around 11.

**Table 7.** Language model performance on Swiss-Prot 2017_03 data. The downstream classification task is level 1 EC prediction on the EC50 dataset as described below.

| Split | # Train | LM |  | Downstream |
| --- | --- | --- | --- | --- |
|  |  | Perpl. | Acc. | Acc. |
| UniRef50 | 316K | 11.37 | 0.254 | 0.956 |
| UniRef50 (clean) | 322K | 11.75 | 0.244 | 0.954 |
| Random(64% train) | 316K | 7.40 | 0.385 | 0.955 |
| Random(fwd) | 499K | **6.88** | **0.409** | 0.960 |

Even more important than the language model performance itself is, however, in our present context its impact on downstream classification tasks. This is illustrated exemplarily for the EC level 1 classification performance on the EC50 dataset as described below. Table 7 lists the downstream performance for all language models pretrained on the respective Swiss-Prot version. Interestingly, all fine-tuned classification models show a comparable performance and it is not possible to discriminate between different language models used for pretraining. The same pattern was observed across all considered classification tasks. In particular, it shows that the data leakage from inconsistent splits between pretraining and classification task is presumably small in the context of language modeling. We also experimented with the use of byte-pair-encoding to form subword units [Sennrich et al., 2015]. The language model metrics are obviously not comparable for different vocabulary sizes, so the downstream classification performance is the only meaningful metric to compare both approaches in this case. However, we did not see any significant performance improvements compared to the baseline models with single amino acids as tokens. In addition, if one tokenizes using subword units, sequence annotation tasks such as secondary structure prediction are less straightforward to handle.

Key insights from results presented in this section are the following: First, the results demonstrate that language modeling on amino acid sequence data is indeed meaningful and represents a potentially interesting domain of application for language model methods from NLP. Prediction accuracies of 40% (random split) or 25% (UniRef50 clusters) document a solid understanding of the general structure of proteins. However, the section also highlights crucial differences compared to language model evaluation in NLP. Whereas in NLP sequence similarity between train and test sequences is barely considered, it has crucial implications in proteomics. Here, it is the dataset coverage in terms of clusters rather than the nominal size of the training set, which most crucially determines the language model performance. Second, however, the different language models show no significant differences in terms of downstream classification performance. This iterates a general insight from NLP in the sense that improved language model performance does not necessarily imply improved downstream task performance.

## 4 Evaluation Procedure for Comparison to DEEPre and ECPred

For *DEEPre* the reported accuracies derived from a 5-fold-cross-validation are straightforward to compute. For *ECPred*, however, we reproduced the evaluation scheme of the authors, which is tailored to binary one-vs-all-classifiers opposed to multi-class-classifiers. The scheme requires to evaluate binary classifiers that distinguish a particular EC class from other EC classes as well as non-enzymes. Finally, the mean *F*_1_-score is reported across the six datasets. We adopt their evaluation procedure in order to be able to compare directly to their reported results. However, we decided to fit a seven-class categorical classifier (non-enzymes and six main enzyme classes) both on the concatenated training set for all EC classes. In our experiments, the performance of these classifiers was comparable or even better than the corresponding score obtained by training six independent classifiers on EC-class-specific training sets and the procedure is more in line with our approach. We would like to add that this statement applies only to level 0 and level 1, whereas hierarchical classifiers as used by *ECPred* show advantages for sparsely populated cases such as level 2 EC prediction, which is, however, not considered here. At this point we would like to stress that the evaluation procedure is tailored specifically to hierarchical classifiers rather inconvenient to apply for multi-class classifiers.

## 5 Dependence of Data Leakage on BLAST Database Size

In this section, we investigate the data leakage effect discussed in Section 3.2.2 and Appendix 3 in more detail with focus on PSSM features. The aim is to illustrate that the overstimation of the model generalization performance by computing PSSM features on the whole dataset is not only a theoretical issue but has practical implication for downstream classification performance at least in the cases where the corresponding BLAST database is small.

For pretraining using language modeling this issue did not have significant impact on the downstream performance. This is obviously a desirable property as it would otherwise require to pretrain on large datasets with train-test splits consistent with the downstream task, which defeats the purpose of using pretraining as a universal step unspecific to the choice of the downstream task. However, data leakage is always a potential issue in this context and deserves further research from our perspective to understand the practical implications for different classification tasks.

To highlight the importance of using an appropriate train-test-split also for the database on which PSI-BLAST is performed, we conducted the following experiment, where we compared two sets of features (as already described in Section 3.2.2) both trained with the same model and hyperparameters: (1) PSSM features based on the whole Swiss-Prot database (including test sequences) and (2) PSSM features on database consisting only of sequences from the training clusters. While experimenting, we observed that the effect is consistent but barely measurable for large training data, but as the training data (and therefore the BLAST database) get smaller, the effect becomes more apparent. We considered four training sizes: 10%, 30%, 50% and 80% training size, while the test size stayed 20% (the rest of the sequences were neglected for this experiment. For each experiment, we trained a model for level 1 EC prediction as described in Section 2.2 and shown in Section 3.2.2, where we used an early stopping criterion based on the accuracy on a validation set. Here, we used the EC40 dataset as described in Table 5.

Figure 3 shows the results of this experiment. It can be seen that the effect (difference between clean and leakage) is getting more pronounced when the training dataset gets smaller. This might be a obvious fact from a machine learning point of view, but to the best of our knowledge, this has never been shown in the context of PSSM features for proteomics. From this finding, we can conclude that the results from previous literature are not overestimated strongly. Nevertheless, a fundamental difference remains when dealing with smaller datasets. In the case of small data, considering e.g. 10% of the EC40 training data, the gap in performance between a clean and a leaky procedure is 8% in accuracy, which is far from negligible. In addition, we restricted our analysis to EC classification whereas the effect might very well also depend on the considered downstream task.

**Figure 3:**
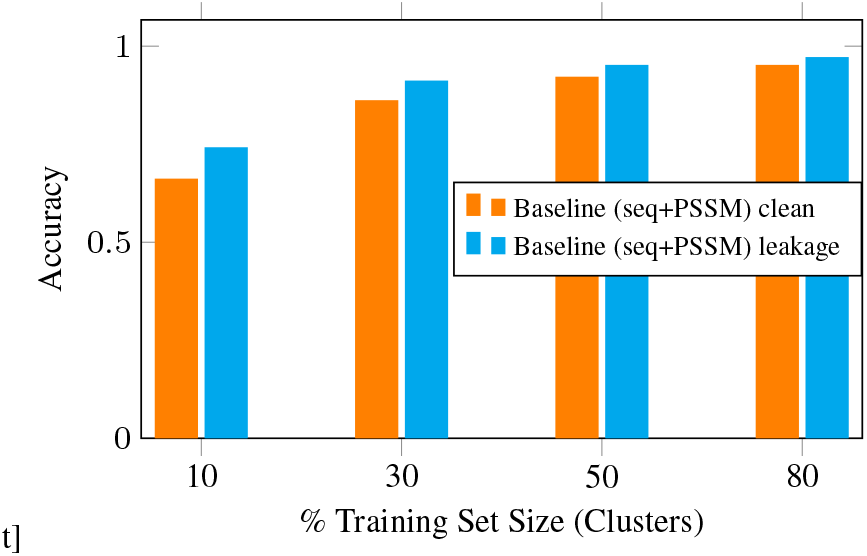
Dependence of EC classification accuracy on the size of the dataset used to computed PSSM features compared to a model exploiting PSSM features computed on the full dataset.

